# Myositis-specific autoantibodies recognizing Mi2 also target the autoimmune regulator (AIRE) protein at a shared PHD-zinc finger

**DOI:** 10.1101/2025.01.15.633218

**Authors:** Jon Musai, Sahana Jayaraman, Katherine Pak, Iago Pinal-Fernandez, Sandra Muñoz-Braceras, Maria Casal-Dominguez, Eric Cho, Fa’alataitaua M. Fitisemanu, Peter D. Burbelo, Mariana J. Kaplan, Blake M. Warner, Adam I. Schiffenbauer, Albert Selva-O’Callaghan, José César Milisenda, Lisa G. Rider, H. Benjamin Larman, Andrew L. Mammen

**Author notes:** **Address correspondence to**: Andrew L. Mammen, M.D., Ph.D., or H. Benjamin Larman, Ph.D, Muscle Disease Section, National Institute of Arthritis and Musculoskeletal and Skin Diseases, National Institutes of Health, 50 South Drive, Room 1141, Building 50, MSC 8024, Bethesda, MD 20892. or. These authors contributed equally (Jon Musai, Sahana Jayaraman, H. Benjamin Larman, Andrew L. Mammen).

## Abstract

**Objectives:** In dermatomyositis patients with anti-Mi2 autoantibodies, autoantibodies can enter muscle cells, leading to the aberrant expression of genes normally repressed by the Mi2/nucleosome remodeling and deacetylation (NuRD) complex. However, the mechanism by which autoantibodies interfere with Mi2/NuRD function remains unclear. This study aimed to identify additional autoantibodies in anti-Mi2-positive patients as well as the specific epitopes recognized by anti-Mi2 and any novel autoantibodies.

**Methods:** Phage ImmunoPrecipitation Sequencing (PhIP-Seq) was used to screen serum samples from anti-Mi2-positive myositis patients for autoantibodies. Enzyme-linked immunosorbent assays (ELISA) and luciferase immunoprecipitation system (LIPS) immunoassays were used to detect autoantibodies in serum samples from myositis patients and healthy controls.

**Results:** PhIP-Seq identified autoantibodies recognizing the autoimmune regulator (AIRE) in sera from anti-Mi2 autoantibody-positive patients. Both anti-AIRE and anti-Mi2 autoantibodies predominantly recognized a homologous region of the plant homeodomain zinc finger type I (PHD1), which is critical for AIRE and Mi2/NuRD function. ELISA and LIPS testing showed that anti-Mi2 autoantibody-positive patients were positive for anti-AIRE autoantibodies, while AIRE reactivity was largely absent in healthy comparators, anti-Mi2 autoantibody-negative-myositis, and other autoimmune diseases. Affinity-purified anti-Mi2 autoantibodies recognized both Mi2 and AIRE by ELISA, whereas anti-Mi2-depleted immunoglobulin fractions did not recognize either protein.

**Conclusions:** Autoantibodies recognizing Mi2 also recognize AIRE at a homologous PHD1 finger. This region is required by the Mi2/NuRD complex to anchor the nucleosome and consequently repress gene expression. Our findings suggest that anti-Mi2 autoantibodies disrupt NuRD complex function by binding to the PHD1 domain. Further studies are needed to determine if anti-Mi2 autoantibodies bind other PHD1-containing proteins and their functional implications.

## INTRODUCTION

Myositis is a heterogenous family of autoimmune diseases that includes dermatomyositis (DM), the antisynthetase syndrome (ASyS), immune-mediated necrotizing myopathy (IMNM), and inclusion body myositis (IBM) [1, 2]. Approximately 70% of DM patients have a known myositis-specific autoantibody (MSA) targeting one or more intracellular autoantigens. The most prevalent MSAs in DM target the chromodomain-helicase-DNA binding proteins (CHD3 – i.e., Mi2α or CHD4 – i.e., Mi2β), nuclear matrix protein (NXP2), transcriptional intermediary factor 1 gamma (TIF1ꝩ), or melanoma differentiation-associated protein 5 (MDA5) [1]. Importantly, each of these autoantibodies is associated with a unique disease phenotype. For instance, adult patients with anti-TIF1ꝩ autoantibodies exhibit a significantly heightened risk of malignancy compared to other DM patients but have a lower propensity for developing interstitial lung disease [3]. Conversely, those harboring anti-MDA5 autoantibodies are prone to rapidly progressive interstitial lung disease, with a relatively low risk of cancer [4].

Compared to other patients with DM, those with anti-Mi2 autoantibodies have more severe muscle involvement, more extensive myofiber necrosis, higher muscle enzyme levels, as well as photosensitive rashes [5]. We recently showed that anti-Mi2 autoantibodies can accumulate in the nuclei of muscle cells where they disrupt the function of the Mi2/NuRD complex and induce the overexpression of genes that this protein complex normally represses. [6–8] However, the mechanism by which anti-Mi2 autoantibodies disrupt the function of Mi2 has remained unclear.

The current study initially aimed to identify novel myositis autoantibodies. We performed PhIP-Seq with an oligonucleotide library encoding 274,207 overlapping peptides spanning the human proteome to screen patient sera for novel myositis autoantibodies [9, 10]. Surprisingly, this revealed that sera from individuals with anti-Mi2 autoantibodies also recognize autoimmune regulator (AIRE), a protein whose dysfunction leads to autoimmunity. Anti-Mi2/AIRE autoantibodies recognize a homologous PHD zinc finger that is critical for the function of both Mi2 and AIRE. In the case of Mi2, this domain is required for the Mi2/NuRD complex to interact with nucleosomes. Our findings suggest that anti-Mi2 autoantibodies bind to this domain, prevent the Mi2/NuRD complex from interacting with nucleosomes, and cause the derepression of genes that are normally repressed by this complex.

## METHODS

### Patients and Serum Samples

To conduct PhIP-Seq, we established a discovery cohort comprising 10 anti-Mi2 autoantibody-positive myositis sera: 8 from patients with adult DM and 2 from patients with juvenile DM. Additionally, we utilized 804 serum samples from healthy controls obtained from the Vaccine Research Cohort [11, 12].

To validate our PhIP-Seq findings, we first conducted enzyme-linked immunosorbent assays (ELISAs) to measure anti-AIRE and anti-Mi2β levels in a separate cohort we termed the “validation” cohort. This validation cohort included 26 anti-Mi2 autoantibody-positive patients – 3 of whom were present in the PhIP-Seq discovery cohort. Our validation cohort also included 63 healthy controls and 44 patients with other myositis-specific autoantibodies (Table 1).

**Table 1:**
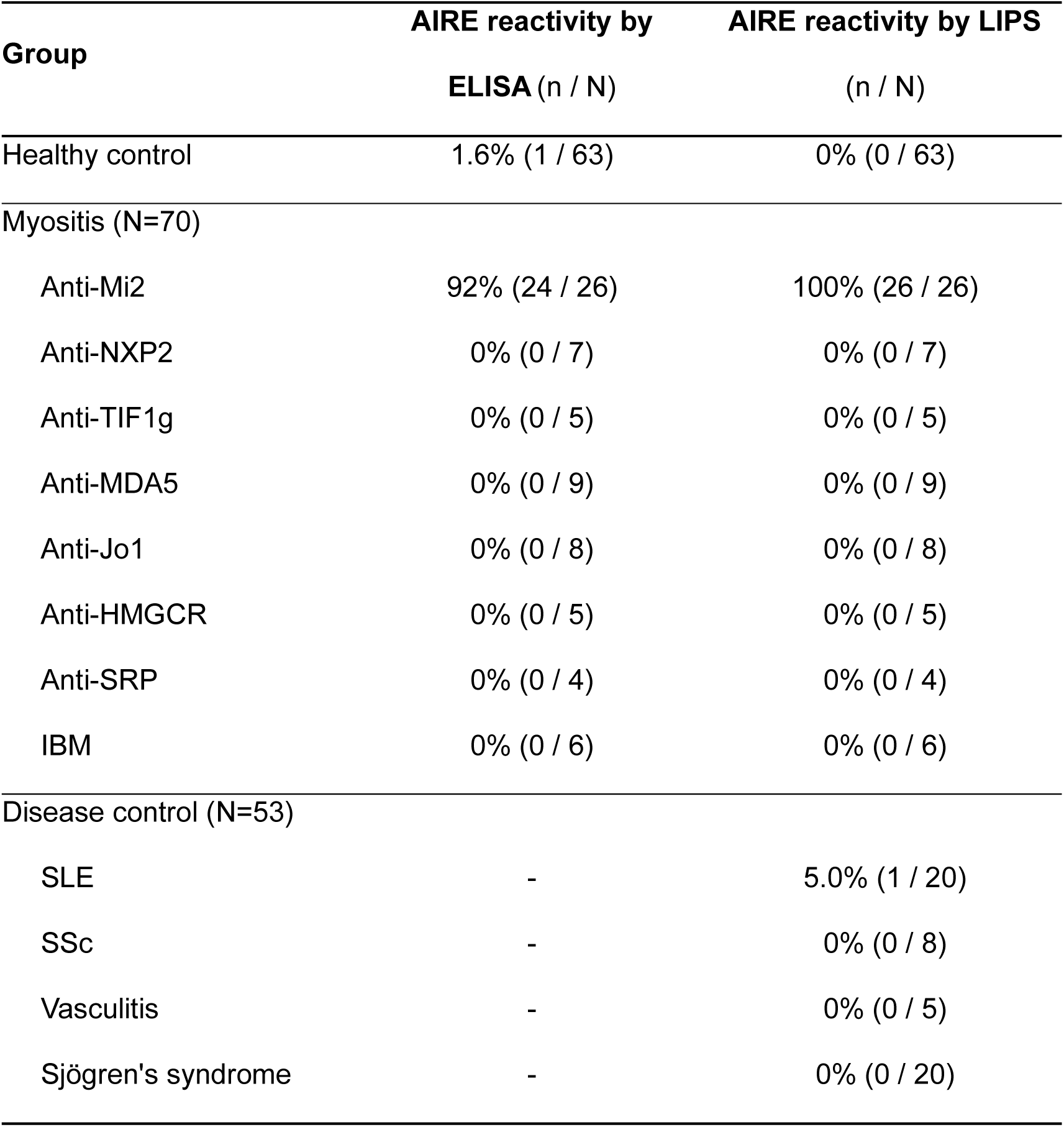
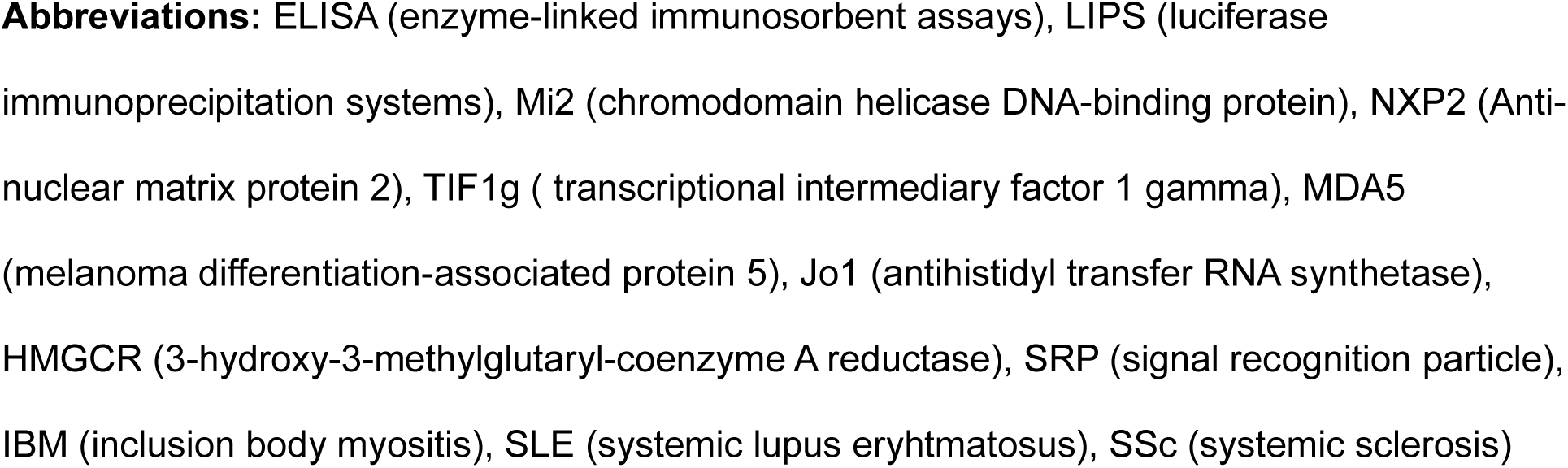
Serum samples tested for anti-AIRE autoantibodies using ELISA and LIPS. Using enzyme-linked immunosorbent assays (ELISAs) and luciferase immunoprecipitation systems (LIPS), the prevalence of anti-AIRE autoantibodies was measured in a validation cohort that included different types of myositis and other healthy and disease comparators.

Since anti-Mi2 autoantibodies can target different chromodomain-helicase-DNA-binding proteins (CHD), like Mi2β (CHD4) or Mi2α (CHD3), we also used luciferase immunoprecipitation system (LIPS) immunoassays to measure anti-AIRE and anti-Mi2α levels in the same validation cohort used for the ELISAs. Additionally, LIPS immunoassays were conducted on 53 serum samples from patients with systemic autoimmune diseases, including 20 with systemic lupus erythematosus (SLE), 8 with systemic sclerosis (SSc), 5 with vasculitis, and 20 with Sjögren’s syndrome (Table 1).

Myositis serum samples were obtained from patients who either fulfilled Llyod’s criteria for inclusion body myositis (IBM) [13] or the Casal and Pinal criteria for other types of autoantibody-positive myositis [1]. Patients were classified as autoantibody-positive if they tested positive for autoantibodies against Mi2, TIF1, NXP2, MDA5, Jo1, HMGCR, or SRP using at least one of the following immunological methods: ELISA, in vitro transcription and translation followed by immunoprecipitation, line blotting (EUROLINE Autoimmune Inflammatory Myopathies 16 Ag (IgG) test kit), or immunoprecipitation from ^35^S-methionine-labeled HeLa cell lysates. All serum samples were obtained from patients enrolled in institutional review board-approved (IRB) cohorts from the NIH in Bethesda, MD as well as the Clinic and Vall d’Hebron Hospitals in Barcelona, Spain.

### Myositis-Specific Autoantibody Discovery and Epitope Mapping by Phage ImmunoPrecipitation Sequencing (PhIP-Seq)

The standard PhIP-Seq procedure [9, 10] was employed to screen sera from 10 anti-Mi2 autoantibody-positive myositis patients. The IgG concentration of each serum sample was quantified through an ELISA, enabling the standardization of IgG input (2 μg per reaction) into the subsequent PhIP-Seq assay. Serum antibodies were incubated overnight with the human peptidome library consisting of 274,207 peptides of 90 amino acids in length. Following incubation, antibody and antibody-bound T7 phage were isolated using protein A and protein G coated Dynal beads (#10002D and #10004D, Invitrogen). The immunoprecipitated phage library underwent PCR amplification with sample-specific DNA barcodes and the resultant amplicons were pooled and sequenced using an Illumina NextSeq instrument. Samples were then demultiplexed and aligned through an informatics pipeline as previously described [11, 14].

### Anti-AIRE and anti-Mi2β ELISA

ELISA plates (#351172, Falcon) were pre-coated overnight at 4°C with 100 ng of recombinant protein of human AIRE (#TP313497M, OriGene) per well diluted in 100 uL of 1X phosphate-buffered saline (PBS). After coating, ELISA plates were washed with PBS-0.05% Tween (PBS-T) and blocked with 300 uL of 5% bovine serum albumin (BSA) in PBS-T for 1 hour at 37°C. Plates were then washed again with PBS-T. Diluted human serum samples (100 uL, 1:400 in 1% BSA/PBS-T) were added to each well and incubated for 1 hour at 37°C and then washed with PBS-T. Diluted HRP-labelled goat anti-human IgG antibody (100 uL, 1:10,000 in 1% BSA/PBST; #109-036-088, Jackson ImmunoResearch Lab) was added and incubated for 30 minutes at 37°C. After washing the plate with PBS-T followed by PBS, 100 uL of Sure Blue Peroxidase Substrate (#52-00-03, KPL) was added. Reactions were stopped with 100 uL of 1N hydrochloric acid. The absorbance at 450 nm was determined, and test sample absorbances were normalized to the sera of an arbitrary positive control sample used as a reference in the anti-AIRE ELISA. The cut-off for determining AIRE-positivity was defined as two standard deviations above the mean normalized absorbance of healthy control sera. We followed this procedure to develop an anti-Mi2β ELISA, adapting by pre-coating with 50 ng of Mi2β protein (#PRO-112, ProSpec) per well.

### Anti-AIRE and anti-Mi2α LIPS

Luciferase immunoprecipitation systems (LIPS) immunoassays were performed as described elsewhere [15–18]. AIRE or Mi2α autoantigen were fused to a light-emitting luciferase and then immunoprecipitated. Synthetic DNA encoding AIRE (NP_000374.1, amino acids 2-545) and Mi2α (NP_005843.2, amino acids 2-596) were obtained from Twist Biosciences. These DNAs contained *BamH1* and *Xho1* restriction sites for directional cloning into the pREN2 eukaryotic expression vector as a C-terminal *Renilla* luciferase fusion protein. Both pREN2-AIRE and - Mi2α constructs were sequenced to verify their integrity. These mammalian expression vectors were then transfected into Cos1 cells using Lipofectamine 3000. Crude lysates were harvested 48 hours after transfection for AIRE and 24 hours for Mi2α. For testing, serum samples from myositis patients, healthy controls, and other autoimmune disease subjects were diluted in assay buffer A (20 mM Tris, pH 7.5 150 mM NaCl, 5 mM MgCl_2_, 1% Triton X-100) and incubated in a 96-well microtiter plate for 1 hour with 1 × 10^7^ light units (LU) per well for either *Renilla* luciferase-Aire or -Mi2α. Next, the serum-antigen mixture was transferred to a microtiter filter plate (Millipore) containing protein Ultralink protein A/G beads (Invitrogen) and incubated for another hour. The filter plates containing the immune complexes were then washed eight times with buffer A and twice with 1X PBS to remove unbound antigens. After the final wash, coelenterazine substrate (Promega) was added to detect the amount of immunoprecipitated *Renilla* luciferase fusion proteins in light units (LU) using a Berthold LB 960 Centro microplate luminometer (Berthold Technologies, Bad Wildbad). The cut-off for determining AIRE-positivity was defined as the mean value plus five standard deviations of the healthy control sera.

### Purification of myositis-specific autoantibodies from patient serum

Human immunoglobulin G (IgG) was purified and concentrated from anti-Mi2 autoantibody-positive (n=3) and anti-MDA5 autoantibody-positive (n=1) sera using protein G Agarose (#16-266, Millipore) and the Amicon Pro Purification System (#ACS500024, Millipore) with a 30 kDa molecular weight cutoff Amicon Ultra Centrifugal Filter (Millipore, ref. UFC503024). After isolating IgG from anti-Mi2 autoantibody-positive serum samples, we used Mi2β-coated magnetic beads to affinity-purify anti-Mi2β autoantibodies, while also preserving the non-bound fraction for further analysis. NHS-activated magnetic beads (#88826, Thermo Scientific) were coupled with recombinant Mi2β human protein (#PRO-112, ProSpec) during an overnight incubation at 4°C with continuous mixing. The unbound ligand was removed, and the beads were washed with quench buffer (100 mM Tris-HCl, 150 M NaCl, pH 8.0). For affinity purification, total IgG from anti-Mi2-positive sera was added to the Mi2β-coupled beads, followed by incubation with continuous mixing for two hours at room temperature. After incubation, the Mi2β-coupled beads were placed on a magnetic rack to separate them from the solution. The supernatant containing unbound IgGs was collected, while the beads were washed to remove residual impurities. Bound IgGs were then eluted using 0.1M glycine at pH 2.0, with the pH neutralized afterward using 10% 1M Tris at pH 8.0, yielding the Mi2β-enriched fraction. To generate the Mi2-depleted fraction, the collected supernatant was re-incubated with fresh Mi2-coupled beads, and this process was repeated three times. This same procedure was performed on purified IgG from the anti-MDA5 autoantibodypositive serum sample using MDA5 protein (#PRO-1505, ProSpec) -coupled NHS-activated beads to obtain the MDA5-enriched and MDA5-depleted fractions. These fractions served as negative controls for anti-AIRE reactivity. Using both the Mi2β and MDA5 fractions, we then performed anti-Mi2β, anti-human IgG (#359722-004, Thermo Scientific), and anti-AIRE ELISAs to assess immunoreactivity.

### Standard Protocol Approvals and Patient Consents

This study was approved by the Institutional Review Boards (IRB) at the National Institutes of Health, the Vall d’Hebron, and the Clinic Hospitals in Barcelona. Written informed consent was obtained from each participant.

### Statistical analysis

PhIP-Seq analysis between anti-Mi2 autoantibody-positive patients and healthy controls was performed using phipCC as previously described [11]. Dichotomous variables were expressed as percentages and absolute frequencies. Pairwise comparisons for categorical variables between groups were made using Fisher’s exact test using the R programming language. Clustal Omega was used to assess the homology between protein sequences. A 2-sided p-value of 0.05 or less was considered statistically significant with no adjustment for multiple comparisons.

## RESULTS

### Novel autoantibody reactivity against AIRE using PhIP-Seq

Among 10 anti-Mi2 autoantibody-positive myositis samples tested with PhIP-Seq, 6 showed reactivity to AIRE (Figure 1A). This reactivity was specific to the AIRE peptide spanning amino acids 271-360 (Figure 1A). These same 6 patients who were positive for anti-AIRE also reacted to the Mi2β peptide spanning amino acids 361-450 (Figure 1B), and 5 of these patients also showed reactivity to the Mi2α peptide spanning amino acids 347-436 (Figure 1C). PhIP-Seq also detected reactivity to other Mi2 peptides, but these were found in 3 or fewer of the 10 anti-Mi2 autoantibody-positive patients. These immunodominant epitopes identified by PhIP-Seq both in AIRE, Mi2β, and Mi2α corresponded to the plant homeodomain zinc finger type I (PHD1), which plays a crucial role in chromatin recognition and transcriptional regulation [19–22]. The immunodominant PHD1 epitopes of the three proteins revealed a high degree of similarity (Figure 1D).

**Figure 1A:**
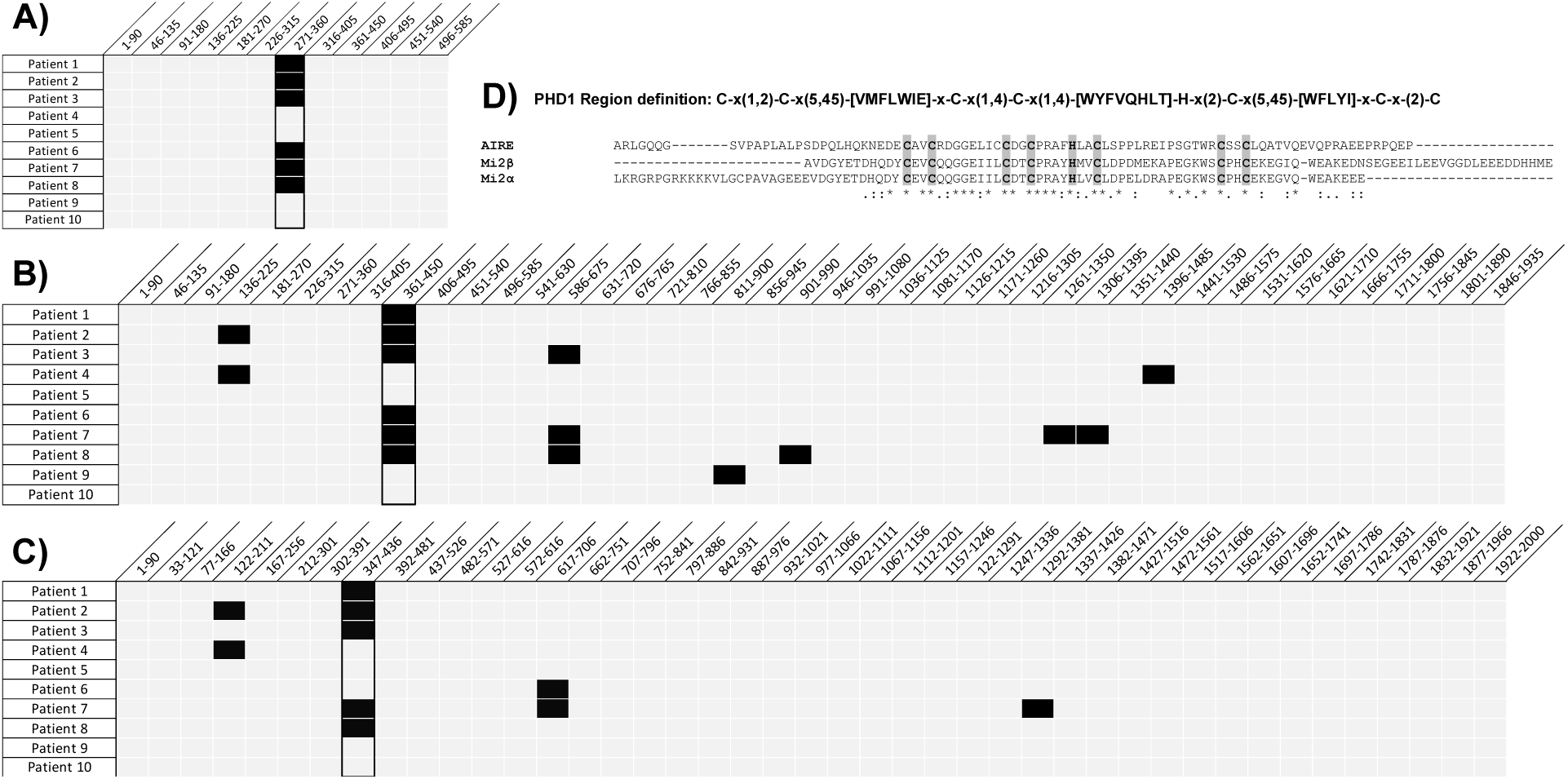
Identification of novel autoantibody specificities through PhIP-Seq analysis. Anti-AIRE (A), anti-Mi2β (B), and anti-Mi2α (C) reactivity as determined by Phage ImmunoPreciptation Sequencing from 10 anti-Mi2 autoantibody-positive myositis patients. For pannels (A) – (C), the peptide fragments of the protein, annotated by their starting and terminal amino acids, are arranged diagonally along the X-axis. Black squares show that the patient has autoantibodies against that peptide fragment, while gray squares indicate that the patient does not have autoantibodies against that peptide. Panel (A) highlights the immunogenic peptides of the AIRE protein, Panel (B) showcases those of the Mi2β protein, and Panel (C) displays the immunogenic peptides of the Mi2α protein. Panel (D) shows sequence alignment of the immunodominant epitopes of AIRE (amino acids 271-360), Mi2β (amino acids 361-450), and Mi2α (amino acids 347-436) using Clustal Omega. The PHD1 region is characterized by a distinct motif featuring a combination of four cysteine residues, one histidine residue, and three additional cysteine residues, which are highlighted in gray. A detailed characterization of the PHD region is provided and can also be found on PROSITE (https://prosite.expasy.org/PS01359). <u>PROSITE (expasy.org)</u>.

### Only myositis patients with anti-Mi2 autoantibodies have anti-AIRE reactivity

We designed an ELISA with full-length AIRE protein to evaluate AIRE reactivity in a validation cohort that consisted of serum samples from 26 anti-Mi2 autoantibody-positive DM patients, 44 myositis patients with other MSAs or IBM, and 63 healthy comparators (Table 1). Among these samples, autoantibodies recognizing AIRE were present in 24/26 (92%) of anti-Mi2 autoantibody-positive and 0/44 (0%) of anti-Mi2 autoantibody-negative myositis patients (Figure 2A). One (1.6%) of the 63 healthy control serum samples had anti-AIRE reactivity (Figure 2A). To validate these data, we used a LIPS immunoassay with an AIRE antigen genetically fused to *Renilla* luciferase to measure autoantibody levels. We tested the same serum samples used in the ELISA, along with samples from 53 patients with different systemic autoimmune diseases. Among these samples, autoantibodies recognizing AIRE were detected in all 26 (100%) of the anti-Mi2 autoantibody-positive patients and none (0%) of the anti-Mi2-negative myositis patients or healthy controls (Figure 2B). Two anti-Mi2 autoantibody-positive samples that were borderline negative for AIRE reactivity by ELISA were identified as positive by LIPS (Figure 2C), likely due to the broader dynamic range of LIPS. Additionally, only 1 (5%) of the 20 SLE sera showed reactivity against AIRE, while none (0%) of the sera from SSc (0/8), vasculitis (0/5), or Sjögren’s syndrome (0/20) patients exhibited AIRE reactivity (Figure 2B). The healthy control with positive anti-AIRE reactivity by ELISA tested negative for AIRE and Mi2 by LIPS and ELISA, suggesting a false positive, and the SLE patient with positive anti-AIRE reactivity showed no anti-Mi2α reactivity by LIPS. We also compared the levels of AIRE-reactive autoantibodies measured by ELISA and LIPS immunoassays, which revealed a strong correlation between the two methods (Figure 2C). This underscores the reliability of both techniques in detecting AIRE reactivity.

**Figure 2:**
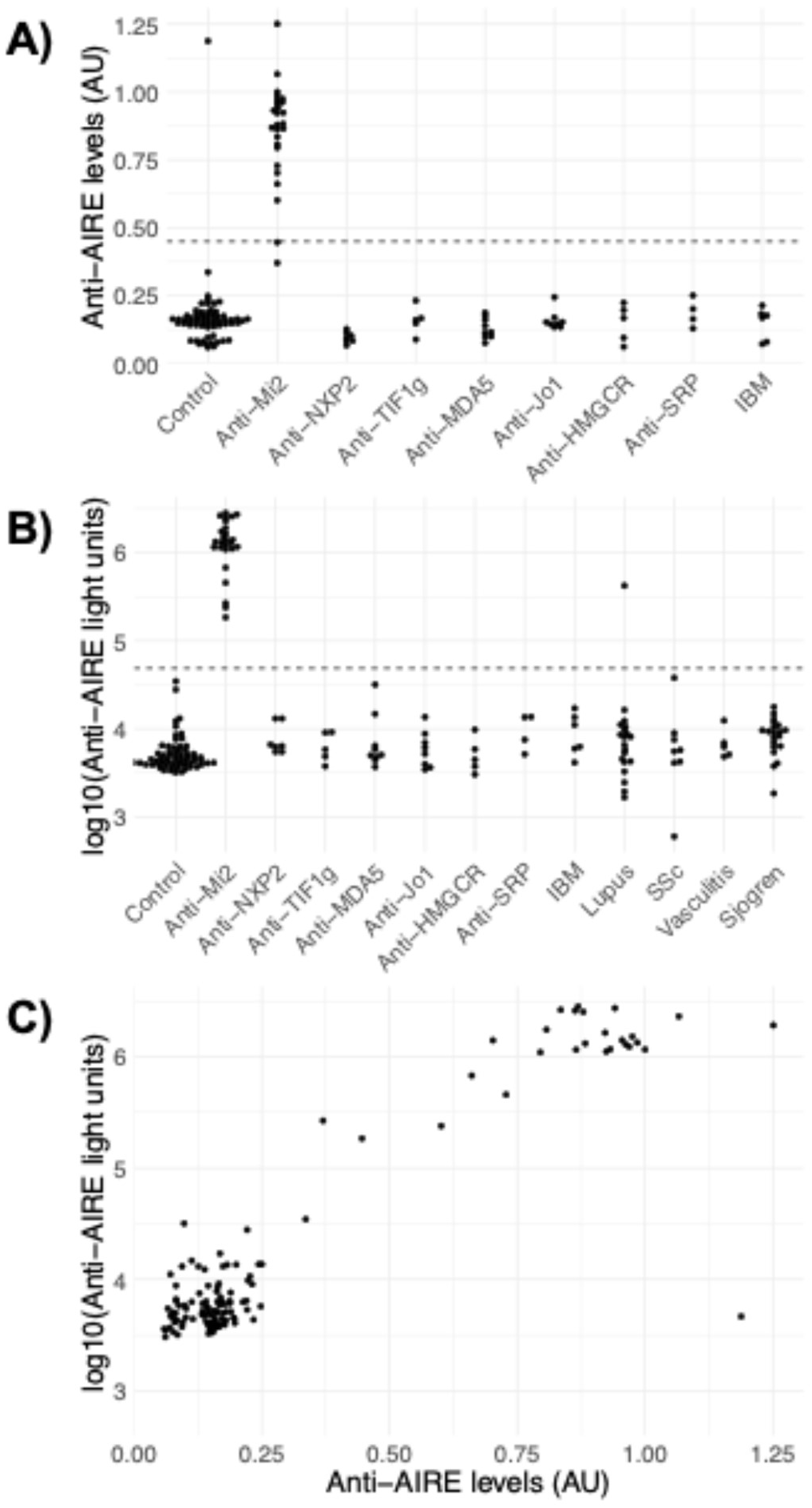
Detection of anti-AIRE autoantibodies in healthy controls, myositis patients, and individuals with other systemic autoimmune diseases. Anti-AIRE autoantibody levels were measured using (A) enzyme-linked immunosorbent assay (ELISA) and (B) luciferase immunoprecipitation system (LIPS) in sera from healthy controls, patients with myositis, and patients with other systemic autoimmune diseases (the latter only for LIPS). The dotted line indicates the cutoff for defining anti-AIRE autoantibody positivity, with the cutoff set at 3 standard deviations for the anti-AIRE ELISA and 5 standard deviations for the anti-AIRE LIPS assay. Panel (C) shows the correlation of anti-AIRE levels measured by ELISA and LIPS in 26 anti-Mi2-positive serum samples, 44 samples with other myositis-specific autoantibodies or inclusion body myositis, and 63 healthy controls by both ELISA (X-axis) and LIPS (Y-axis). AU: arbitrary units.

### Anti-Mi2 autoantibodies also recognize AIRE

Due to the specificity of anti-AIRE reactivity in sera containing anti-Mi2 autoantibodies, and the similarity in the immunodominant epitopes between AIRE and Mi2 proteins, we then explored whether anti-Mi2 autoantibodies target both AIRE and Mi2β. To this end, we affinity-purified anti-Mi2 autoantibodies using full-length Mi2β protein conjugated to magnetic beads from three serum samples that were positive both for anti-Mi2 and anti-AIRE autoantibodies by ELISA. We also collected the non-bound antibodies from this purification process to obtain the anti-Mi2 depleted fraction. We detected AIRE recognition only in the anti-Mi2 enriched fractions whereas depletion of anti-Mi2 autoantibodies resulted in a loss of AIRE reactivity (Figure 3). These results demonstrate that autoantibodies that target Mi2β also recognize AIRE.

**Figure 3A:**
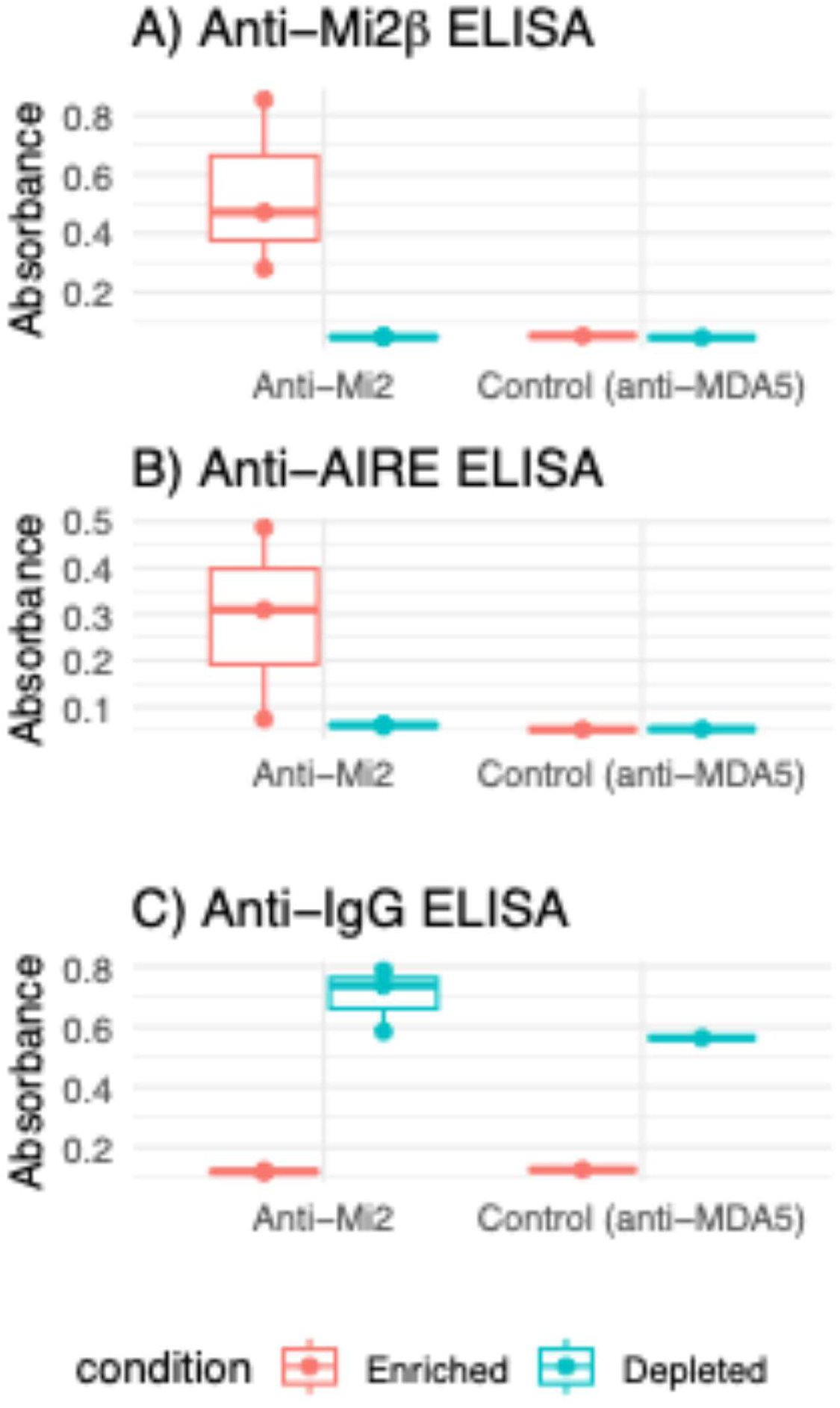
Detection of anti-AIRE reactivity in affinity-purified anti-Mi2 autoantibody fractions. Purified immunoglobulin G (IgG) of anti-Mi2 autoantibody-positive sera (n=3) underwent affinity purification using Mi2β-bound magnetic beads, which produced a Mi2β-enriched and Mi2β-depleted fraction. As a control, purified IgG of one anti-MDA5 autoantibody-positive serum sample underwent affinity purification using MDA5-bound magnetic beads, which produced an MDA5-enriched and MDA5-depleted fraction. These fractions were subjected to an ELISA to detect immunoreactivity against Mi2β (A) and AIRE (B). The total concentration of immunoglobulin was determined by ELISA in the two sets of samples (C).

### Anti-AIRE levels do not correlate with anti-Mi2 levels

Given that nearly all anti-Mi2 autoantibody-positive patients were also anti-AIRE-positive, it was not feasible to conduct a comparative analysis of the clinical characteristics between anti-AIRE-positive and anti-AIRE-negative patients within anti-Mi2 autoantibody-positive myositis. However, given that anti-Mi2 autoantibody levels correlate with disease severity [5, 6], we evaluated the association between anti-AIRE and anti-Mi2β levels within the 26 anti-Mi2 DM/JDM patient sera in our validation cohort. We observed a poor correlation between anti-AIRE and anti-Mi2β levels as determined by ELISA (Figure 4A). As noted earlier, anti-Mi2 autoantibodies can also recognize Mi2α (CHD3). Therefore, we employed LIPS immunoassays to assess whether anti-AIRE levels correlate with anti-Mi2α levels. We also observed a weak correlation between anti-AIRE and anti-Mi2α levels by LIPS (Figure 4B).

**Figure 4:**
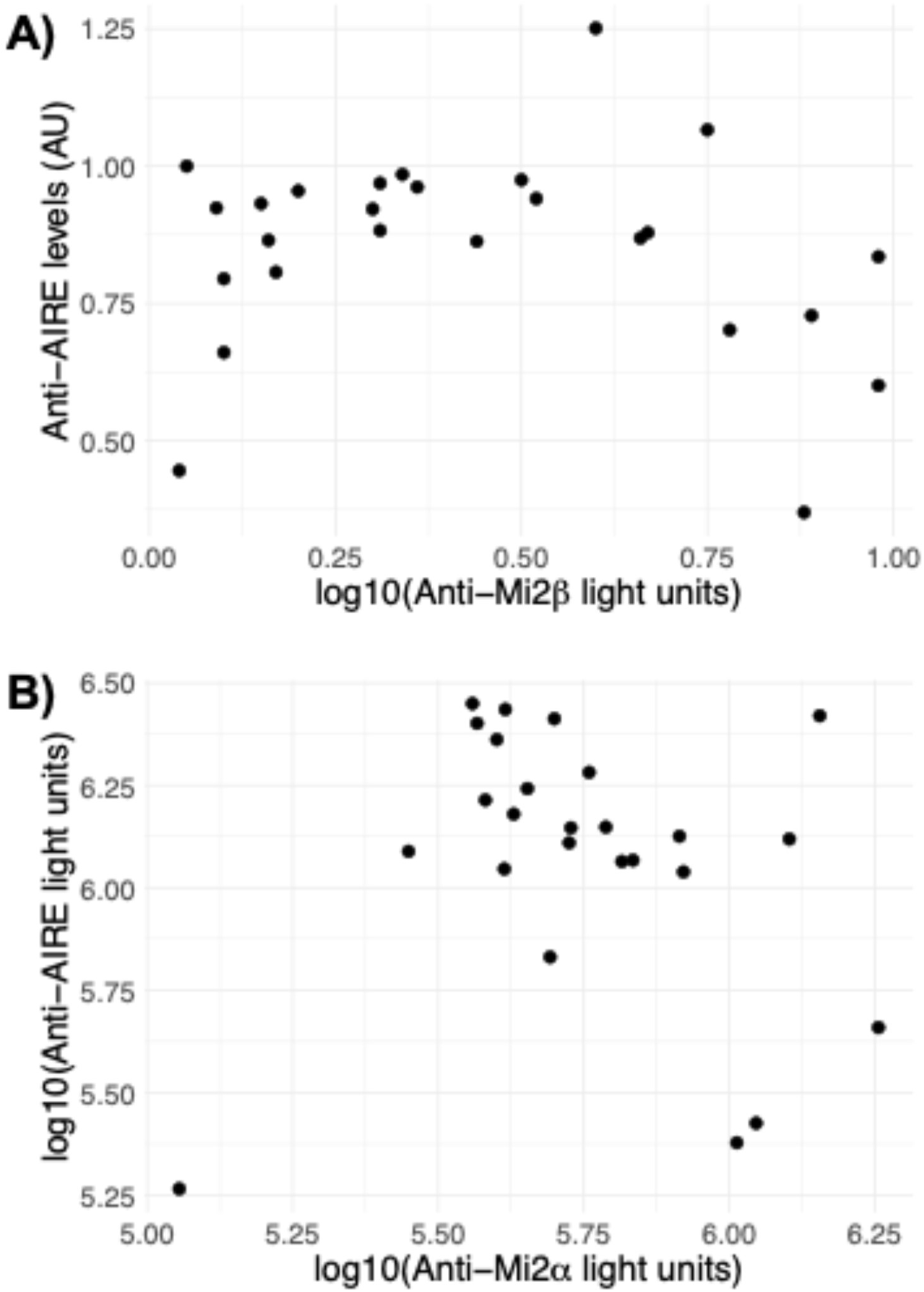
Correlation between anti-AIRE and anti-Mi2 autoantibody reactivity levels by ELISA and LIPS. (A) Reactivity levels measured by ELISA (measured in arbitrary units [AU]) of anti-AIRE and anti-Mi2β autoantibodies and (B) Reactivity levels measured by LIPS (measured in light units) of anti-AIRE and anti-Mi2α autoantibodies. AU: arbitrary units.

## DISCUSSION

In this study, we used PhIP-Seq to identify novel autoantibodies in the serum of DM patients with anti-Mi2 autoantibodies. We found that sera from individuals with autoantibodies against Mi2 also recognize the AIRE protein. Importantly, PhIP-Seq allowed us to identify the PHD1 region as a shared immunodominant epitope of the AIRE and Mi2 autoantigens. This finding is consistent with a previous study defining the immunogenic regions of Mi2 [23]. We also demonstrated that affinity-purified anti-Mi2 autoantibodies can recognize AIRE. When these autoantibodies are removed from the total immunoglobulin fraction, AIRE reactivity is lost. This highlights that the same autoantibodies bind both Mi2 and AIRE, presumably at the PHD1 finger. The PHD1 finger is distinguished by a unique motif consisting of four cysteine residues, one histidine residue, and an additional three cysteine residues. This region is critical for both AIRE and Mi2 to recognize histone modifications and regulate target gene expression by coordinating with transcriptional and chromatin-remodeling machinery [19–22].

AIRE is a pivotal transcriptional regulator that maintains central immune tolerance by controlling the expression of tissue-restricted self-antigens in the thymus [22, 24]. This is crucial for eliminating autoreactive T cells or redirecting them towards a regulatory T cell lineage [22, 24]. AIRE’s role also extends beyond the thymus [25, 26]. Peripheral AIRE is mainly expressed in “extra-thymic AIRE-expressing cells,” which also demonstrate antigen-presenting capabilities and are hypothesized to assist in peripheral immune tolerance [27–34].

Mutations in *AIRE* lead to a rare monogenic disorder called autoimmune polyendocrinopathy-candidiasis-ectodermal dystrophy (APECED), and mono-allelic mutations in the PHD1 finger of AIRE have been found to associate with a broader range of autoimmune phenotypes [35–39]. Moreover, mutations in the PHD1 finger of *AIRE* have been shown to significantly reduce the expression of AIRE-targeted genes and impair AIRE’s binding to their promoters [19, 20]. This highlights the PHD1 finger as a critical domain for AIRE’s function. Myositis-specific autoantibodies targeting the PHD1 region of AIRE could in theory disrupt its role in regulating tissue-restricted antigen expression, potentially leading to immune tolerance breakdown. However, muscle biopsies from anti-Mi2 myositis patients show the transcriptional derepression of genes, including Mi2-regulated genes and others not typically expressed in skeletal muscle [6]. Given that AIRE promotes gene expression, we hypothesize that anti-AIRE autoantibodies, if functionally relevant, would repress gene expression, which contradicts the observed transcriptional profile in anti-Mi2 muscle biopsies. Additionally, although AIRE dysfunction is known to cause autoimmunity, we found no clinical reports indicating APECED-related manifestations in either pediatric [40] or adult [5] populations of anti-Mi2 autoantibody-positive patients. This suggests that the Mi2/NuRD complex is likely the main target of these myositis-specific autoantibodies, through which they exert their pathological effects. Future studies are needed to determine whether these anti-Mi2/AIRE autoantibodies impair AIRE’s function in the thymus or other cells that express this transcriptional regulator, which is crucial for preventing autoimmunity.

Previously, we demonstrated that anti-Mi2 autoantibodies are internalized by muscle cells. They can accumulate in the nucleus and disrupt the function of the Mi2/NuRD complex, which normally represses a specific gene set [6, 7]. Binding of histone 3 tails by PHD fingers is crucial for the gene regulatory function of the Mi2/NuRD complex [41]. Therefore, anti-Mi2 autoantibodies may contribute to myositis pathophysiology by binding the PHD1 finger of the Mi2/NuRD complex and interfering with its ability to bind chromatin. This may result in the subsequent expression of genes typically repressed by this complex in muscle cells.

Of note, detection of anti-Mi2 autoantibodies using standard commercial tests is often unreliable.[42] The discovery that anti-Mi2 autoantibodies also recognize AIRE suggests that testing for anti-AIRE reactivity may provide an improved detection method, especially for patients who test borderline negative in assays utilizing Mi2 protein. The lack of correlation between anti-Mi2 and anti-AIRE antibody levels may result from autoantibodies targeting regions of Mi2 outside the PHD1 finger (Figure 1). Therefore, detecting anti-AIRE autoantibodies in an anti-Mi2 autoantibody-positive patient can confirm that these autoantibodies are targeting the functionally-relevant PHD1 finger of Mi2, as this is the only shared epitope between both proteins. Alternatively, assays specifically designed to target the PHD1 finger may provide a more physiologically-relevant detection compared to currently available methods. However, larger studies, including those addressing false positives in anti-Mi2 results, are needed to robustly support these findings.

In summary, we found that anti-Mi2 autoantibodies also recognize AIRE protein at a shared epitope in the PHD-zinc finger domain of both Mi2 and AIRE proteins. This suggests that anti-Mi2 autoantibodies disrupt the function of the Mi2/NuRD complex by preventing it from interacting with chromatin. Given the key role of the PHD1 finger in regulating global transcriptional factors, further investigation is needed to understand how autoantibodies targeting this region affect proteins like AIRE. Exploring the muscle-specific and systemic effects of this reactivity could reveal how the dysregulation of target autoantigens like Mi2 and AIRE interact to drive the development of myositis and other autoantibody-mediated diseases.

## Contributorship

All authors contributed to the development of the manuscript, including interpretation of results, substantive review of drafts, and approval of the final draft for submission.

## Acknowledgments

None.

## Patient and public involvement

Patients and/or the public were not involved in the design, conduct, reporting, or dissemination plans of this research.

## Data sharing statement

Any anonymized data not published within the article will be shared by request from any investigator.

## References

1. Casal-Dominguez M, Pinal-Fernandez I, Pak K, Huang W, Selva-O’Callaghan A, Albayda J, et al. Performance of the 2017 European Alliance of Associations for Rheumatology/American College of Rheumatology Classification Criteria for Idiopathic Inflammatory Myopathies in Patients With Myositis-Specific Autoantibodies. Arthritis Rheumatol. 2022 Mar; 74(3):508–517.

2. Selva-O’Callaghan A, Pinal-Fernandez I, Trallero-Araguas E, Milisenda JC, Grau-Junyent JM, Mammen AL. Classification and management of adult inflammatory myopathies. Lancet Neurol. 2018 Sep; 17(9):816–828.

3. Trallero-Araguas E, Rodrigo-Pendas JA, Selva-O’Callaghan A, Martinez-Gomez X, Bosch X, Labrador-Horrillo M, et al. Usefulness of anti-p155 autoantibody for diagnosing cancer-associated dermatomyositis: a systematic review and meta-analysis. Arthritis Rheum. 2012 Feb; 64(2):523–532.

4. Sato S, Hoshino K, Satoh T, Fujita T, Kawakami Y, Fujita T, et al. RNA helicase encoded by melanoma differentiation-associated gene 5 is a major autoantigen in patients with clinically amyopathic dermatomyositis: Association with rapidly progressive interstitial lung disease. Arthritis Rheum. 2009 Jul; 60(7):2193–2200.

5. Pinal-Fernandez I, Mecoli CA, Casal-Dominguez M, Pak K, Hosono Y, Huapaya J, et al. More prominent muscle involvement in patients with dermatomyositis with anti-Mi2 autoantibodies. Neurology. 2019 Nov 5; 93(19):e1768–e1777.

6. Pinal-Fernandez I, Milisenda JC, Pak K, Munoz-Braceras S, Casal-Dominguez M, Torres-Ruiz J, et al. Transcriptional derepression of CHD4/NuRD-regulated genes in the muscle of patients with dermatomyositis and anti-Mi2 autoantibodies. Ann Rheum Dis. 2023 Aug; 82(8):1091–1097.

7. Pinal-Fernandez I, Munoz-Braceras S, Casal-Dominguez M, Pak K, Torres-Ruiz J, Musai J, et al. Pathological autoantibody internalisation in myositis. Ann Rheum Dis. 2024 Jun 20.

8. Pinal-Fernandez I, Casal-Dominguez M, Derfoul A, Pak K, Miller FW, Milisenda JC, et al. Machine learning algorithms reveal unique gene expression profiles in muscle biopsies from patients with different types of myositis. Ann Rheum Dis. 2020 Sep; 79(9):1234–1242.

9. Larman HB, Zhao Z, Laserson U, Li MZ, Ciccia A, Gakidis MA, et al. Autoantigen discovery with a synthetic human peptidome. Nat Biotechnol. 2011 May 22; 29(6):535–541.

10. Larman HB, Laserson U, Querol L, Verhaeghen K, Solimini NL, Xu GJ, et al. PhIP-Seq characterization of autoantibodies from patients with multiple sclerosis, type 1 diabetes and rheumatoid arthritis. J Autoimmun. 2013 Jun; 43:1–9.

11. Angkeow JW, Monaco DR, Chen A, Venkataraman T, Jayaraman S, Valencia C, et al. Phage display of environmental protein toxins and virulence factors reveals the prevalence, persistence, and genetics of antibody responses. Immunity. 2022 Jun 14; 55(6):1051–1066 e1054.

12. Venkataraman T, Valencia C, Mangino M, Morgenlander W, Clipman SJ, Liechti T, et al. Analysis of antibody binding specificities in twin and SNP-genotyped cohorts reveals that antiviral antibody epitope selection is a heritable trait. Immunity. 2022 Jan 11; 55(1):174–184 e175.

13. Lloyd TE, Mammen AL, Amato AA, Weiss MD, Needham M, Greenberg SA. Evaluation and construction of diagnostic criteria for inclusion body myositis. Neurology. 2014 Jul 29; 83(5):426–433.

14. Morgenlander WR, Chia WN, Parra B, Monaco DR, Ragan I, Pardo CA, et al. Precision arbovirus serology with a pan-arbovirus peptidome. Nat Commun. 2024 Jul 11; 15(1):5833.

15. Burbelo PD, Ching KH, Klimavicz CM, Iadarola MJ. Antibody profiling by Luciferase Immunoprecipitation Systems (LIPS). J Vis Exp. 2009 Oct 7(32).

16. Burbelo PD, Riedo FX, Morishima C, Rawlings S, Smith D, Das S, et al. Sensitivity in Detection of Antibodies to Nucleocapsid and Spike Proteins of Severe Acute Respiratory Syndrome Coronavirus 2 in Patients With Coronavirus Disease 2019. J Infect Dis. 2020 Jun 29; 222(2):206–213.

17. Burbelo PD, Iadarola MJ, Keller JM, Warner BM. Autoantibodies Targeting Intracellular and Extracellular Proteins in Autoimmunity. Front Immunol. 2021; 12:548469.

18. Burbelo PD, Huapaya JA, Khavandgar Z, Beach M, Pinal-Fernandez I, Mammen AL, et al. Quantification of autoantibodies using a luminescent profiling method in autoimmune interstitial lung disease. Front Immunol. 2024; 15:1462242.

19. Koh AS, Kuo AJ, Park SY, Cheung P, Abramson J, Bua D, et al. Aire employs a histone-binding module to mediate immunological tolerance, linking chromatin regulation with organ-specific autoimmunity. Proc Natl Acad Sci U S A. 2008 Oct 14; 105(41):15878–15883.

20. Org T, Chignola F, Hetenyi C, Gaetani M, Rebane A, Liiv I, et al. The autoimmune regulator PHD finger binds to non-methylated histone H3K4 to activate gene expression. EMBO Rep. 2008 Apr; 9(4):370–376.

21. Zhong Y, Moghaddas Sani H, Paudel BP, Low JKK, Silva APG, Mueller S, et al. The role of auxiliary domains in modulating CHD4 activity suggests mechanistic commonality between enzyme families. Nat Commun. 2022 Dec 6; 13(1):7524.

22. Peterson P, Org T, Rebane A. Transcriptional regulation by AIRE: molecular mechanisms of central tolerance. Nat Rev Immunol. 2008 Dec; 8(12):948–957.

23. Ge Q, Nilasena DS, O’Brien CA, Frank MB, Targoff IN. Molecular analysis of a major antigenic region of the 240-kD protein of Mi-2 autoantigen. J Clin Invest. 1995 Oct; 96(4):1730–1737.

24. Anderson MS, Venanzi ES, Chen Z, Berzins SP, Benoist C, Mathis D. The cellular mechanism of Aire control of T cell tolerance. Immunity. 2005 Aug; 23(2):227–239.

25. Zhao B, Chang L, Fu H, Sun G, Yang W. The Role of Autoimmune Regulator (AIRE) in Peripheral Tolerance. J Immunol Res. 2018; 2018:3930750.

26. Anderson MS, Su MA. AIRE expands: new roles in immune tolerance and beyond. Nat Rev Immunol. 2016 Apr; 16(4):247–258.

27. Huo F, Li D, Zhao B, Luo Y, Zhao B, Zou X, et al. Deficiency of autoimmune regulator impairs the immune tolerance effect of bone marrow-derived dendritic cells in mice. Autoimmunity. 2018 Feb; 51(1):10–17.

28. Gardner JM, Devoss JJ, Friedman RS, Wong DJ, Tan YX, Zhou X, et al. Deletional tolerance mediated by extrathymic Aire-expressing cells. Science. 2008 Aug 8; 321(5890):843–847.

29. Zhu W, Yang W, He Z, Liao X, Wu J, Sun J, et al. Overexpressing autoimmune regulator regulates the expression of toll-like receptors by interacting with their promoters in RAW264.7 cells. Cell Immunol. 2011; 270(2):156–163.

30. Lindh E, Lind SM, Lindmark E, Hassler S, Perheentupa J, Peltonen L, et al. AIRE regulates T-cell-independent B-cell responses through BAFF. Proc Natl Acad Sci U S A. 2008 Nov 25; 105(47):18466–18471.

31. Lindmark E, Chen Y, Georgoudaki AM, Dudziak D, Lindh E, Adams WC, et al. AIRE expressing marginal zone dendritic cells balances adaptive immunity and T-follicular helper cell recruitment. J Autoimmun. 2013 May; 42:62–70.

32. Yamano T, Dobes J, Voboril M, Steinert M, Brabec T, Zietara N, et al. Aire-expressing ILC3-like cells in the lymph node display potent APC features. J Exp Med. 2019 May 6; 216(5):1027–1037.

33. Dobes J, Ben-Nun O, Binyamin A, Stoler-Barak L, Oftedal BE, Goldfarb Y, et al. Extrathymic expression of Aire controls the induction of effective T(H)17 cell-mediated immune response to Candida albicans. Nat Immunol. 2022 Jul; 23(7):1098–1108.

34. Lee JW, Epardaud M, Sun J, Becker JE, Cheng AC, Yonekura AR, et al. Peripheral antigen display by lymph node stroma promotes T cell tolerance to intestinal self. Nat Immunol. 2007 Feb; 8(2):181–190.

35. Peterson P, Nagamine K, Scott H, Heino M, Kudoh J, Shimizu N, et al. APECED: a monogenic autoimmune disease providing new clues to self-tolerance. Immunol Today. 1998 Sep; 19(9):384–386.

36. Saugier-Veber P, Drouot N, Wolf LM, Kuhn JM, Frebourg T, Lefebvre H. Identification of a novel mutation in the autoimmune regulator (AIRE-1) gene in a French family with autoimmune polyendocrinopathy-candidiasis-ectodermal dystrophy. Eur J Endocrinol. 2001 Apr; 144(4):347–351.

37. Oftedal BE, Hellesen A, Erichsen MM, Bratland E, Vardi A, Perheentupa J, et al. Dominant Mutations in the Autoimmune Regulator AIRE Are Associated with Common Organ-Specific Autoimmune Diseases. Immunity. 2015 Jun 16; 42(6):1185–1196.

38. Halonen M, Kangas H, Ruppell T, Ilmarinen T, Ollila J, Kolmer M, et al. APECED-causing mutations in AIRE reveal the functional domains of the protein. Hum Mutat. 2004 Mar; 23(3):245–257.

39. Finnish-German AC. An autoimmune disease, APECED, caused by mutations in a novel gene featuring two PHD-type zinc-finger domains. Nat Genet. 1997 Dec; 17(4):399–403.

40. Rider LG, Shah M, Mamyrova G, Huber AM, Rice MM, Targoff IN, et al. The myositis autoantibody phenotypes of the juvenile idiopathic inflammatory myopathies. Medicine (Baltimore). 2013 Jul; 92(4):223–243.

41. Musselman CA, Ramirez J, Sims JK, Mansfield RE, Oliver SS, Denu JM, et al. Bivalent recognition of nucleosomes by the tandem PHD fingers of the CHD4 ATPase is required for CHD4-mediated repression. Proc Natl Acad Sci U S A. 2012 Jan 17; 109(3):787–792.

42. Pinal-Fernandez I, Pak K, Casal-Dominguez M, Hosono Y, Mecoli C, Christopher-Stine L, et al. Validation of anti-Mi2 autoantibody testing by line blot. Autoimmun Rev. 2020 Jan; 19(1):102425.

